# Genetic Algorithms for model refinement and rule discovery in a high-dimensional agent-based model of inflammation

**DOI:** 10.1101/790394

**Authors:** R Chase Cockrell, Gary An

## Abstract

**Introduction:** Agent-based modeling frequently used modeling method for multi-scale mechanistic modeling. However, the same properties that make agent-based models (ABMs) well suited to representing biological systems also present significant challenges with respect to their construction and calibration, particularly with respect to the selection of potential mechanistic rules and the large number of free parameters often present in these models. We have proposed that various machine learning approaches (such as genetic algorithms (GAs)) can be used to more effectively and efficiently deal with rule selection and parameter space characterization; the current work applies GAs to the challenge of calibrating a complex ABM to a specific data set, while preserving biological heterogeneity.

**Methods:** This project uses a GA to augment the rule-set for a previously validated ABM of acute systemic inflammation, the Innate Immune Response ABM (IIRABM) to clinical time series data of systemic cytokine levels from a population of burn patients. The genome for the GA is a vector generated from the IIRABM’s Model Rule Matrix (MRM), which is a matrix representation of not only the constants/parameters associated with the IIRABM’s cytokine interaction rules, but also the existence of rules themselves. Capturing heterogeneity is accomplished by a fitness function that incorporates the sample value range (“error bars”) of the clinical data.

**Results:** The GA-enabled parameter space exploration resulted in a set of putative MRM rules and associated parameterizations which closely match the cytokine time course data used to design the fitness function. The number of non-zero elements in the MRM increases significantly as the model parameterizations evolve towards a fitness function minimum, transitioning from a sparse to a dense matrix. This results in a model structure that more closely resembles (at a superficial level) the structure of data generated by a standard differential gene expression experimental study.

**Conclusion:** We present an HPC-enabled evolutionary computing approach to calibrate a complex ABM to clinical data while preserving biological heterogeneity. The integration of machine learning, HPC, and multi-scale mechanistic modeling provides a pathway forward to effectively represent the heterogeneity of clinical populations and their data.

**Author Summary:** In this work, we utilize genetic algorithms (GA) to operate on the internal rule set of a computational of the human immune response to injury, the Innate Immune Response Agent-Based Model (IIRABM), such that it is iteratively refined to generate cytokine time series that closely match what is seen in a clinical cohort of burn patients. At the termination of the GA, there exists an ensemble of candidate model rule-sets/parameterizations which are validated by the experimental data;

## Introduction

Agent-based modeling is an object-oriented, discrete-event, rule-based, spatially-explicit, stochastic modeling method. In an agent-based model (ABM), individual agents are simulated interacting with each other and with their environment. These interactions are mediated by a pre-defined set of rules. Essentially, ABMs are computational instantiations of mechanistic knowledge regarding the systems being modeled and consequently are often used to simulate complex systems, in which the aggregate of individual agent interactions can lead to non-trivial or unintuitive macro-state/system-level behaviors. This makes agent-based modeling a powerful technique for representing biological systems; rules are derived from experimentally observed biological behaviors, and the spatially-explicit nature of the models give it an inherent ability to capture space/geometry/structure of biological tissue, which facilitates the ability of biomedical researchers to express and represent their hypotheses in an ABM(1). ABM’s have been used to study and model a wide variety of biological systems (2), from general purpose anatomic/cell-for-cell representations of organ systems capable of reproducing multiple independent phenomena (3, 4) to platforms for drug development (5, 6), and are frequently used to model non-linear dynamical systems such as the human immune system (7–10). In this work, we describe how the use of genetic algorithms (GA) can be used to increase the fidelity and utility of ABMs by facilitating the challenge of calibration to heterogeneous clinical data and providing a means of enhancing the interaction rules beyond the necessary initial abstractions in the base ABM.

In the process of developing an agent-based model, hypotheses or pieces of existing knowledge are re-framed as *rules* that determine the behavior of the agents when they interact with each and their environment. For example, in the context of a biomedical ABM one of those rules might be the definition of a cytokine signaling pathway, i.e., Tumor Necrosis Factor α (TNFα) upregulates Interleukin-10 (IL-10). The quantification of the effect that TNFα has on IL-10 in this hypothetical rule is determined during the model calibration, a critical step in the development and refinement of an ABM (11–15).

In our computational models, the rules and a set of coefficients that quantify the effect of the rules are stored in an object which we refer to as the Model Rule Matrix (MRM).In this scheme, specific rules are represented by rows in the matrix; each computationally relevant entity in the model is then represented by the matrix columns. As a simple example, the system of model rule equations for a single cell:

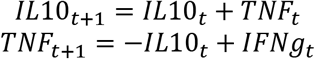

Would be represented by the matrix:

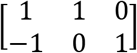

Where the first column is the IL-10 column, the second column is the TNFα column, and the third column is the interferon-gamma (IFN-γ) (another cytokine) column. We note that this is a simplified rule for illustration. The matrix is readily decomposable into a one-dimensional vector, upon which we can operate using genetic algorithms. The genome vector is then padded with an additional three parameters which govern the nature of the injury and how quickly damage spreads though tissue. This addition describes the component of the time evolution of the spatial distribution of a tissue injury that is independent of cytokine levels. The number of rows in the matrix then is equal to the number of rules that it represents, and the number of columns is equal to the number of entities that could potentially contribute to the decision made by their associated rule.

Traditionally, mathematical biological modeling has attempted to ignore (i.e., wash away, integrate out, smooth) variance (which is often sloppily referred to as “noise”) in the data set being modeled (16). As part of the calibration process, a parameter sensitivity analysis (17–19), in which the dependence of mode output on variance in a single parameter (or potentially a combination or parameters) is quantified, is performed. In a complex ABM, this process requires the consideration of a large number of potentially free parameters, making a comprehensive calibration difficult (20–24) and significantly diminishing the utility of traditional parameter sensitivity analysis techniques (25, 26). These difficulties are compounded when considering the range of biological heterogeneity seen experimentally and clinically (9, 27). Additionally, this interpretation disregards the fact that *distinct and unique* genetic “parameterizations” can all present biologically plausible/viable solutions to the problem they are trying to solve – in this model, that is healing from a severe burn. We posit that the aforementioned traditional techniques actively select against a model (and its putative parameterizations) able to capture the biological heterogeneity that is the hallmark of real world clinical scenarios, and that this inability to identify heterogeneity-reproducing models results in brittle models and limits their utility in answering clinically relevant problems (27).

Hence, while sensitivity analysis can be effective at characterizing certain dynamic processes of ABMs, traditional sensitivity analysis applied to the high-dimensional, complex parameter spaces seen in detailed ABMs restricts the representational richness of these models and limits their ability to effectively reproduce and predict the heterogeneous range of real-world data. As an illustrative example, consider the following rule, present in the original version of the Innate Immune Response Agent-Based Model (IIRABM), describing the activation state of a single monocyte:

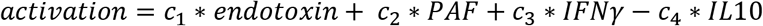

Where the *c*_*i*_^’^*s* represent some constant values used to weight the individual contributions of the array of cytokines. One could perform a sensitivity analysis on a model consisting solely of this rule, however that analysis would miss import bio-plausible model calibrations/parameterizations. While the rule is coded this way in the model, this representation does not actually represent the assertion made by the rule. A more complete way to write this rule would be:

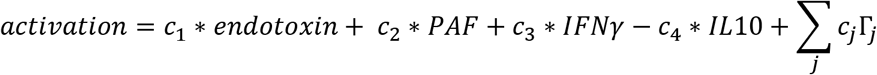

Where *j* sums over the complete set of cytokines on the model, Γ_*j*_ represents the concentration of a specific cytokine, and the constant weighting terms, *c*_*j*_ = 0. Thus, a comprehensive sensitivity analysis must consider implicit zeros in model rule construction, which can vastly increase the size of the task.

Additionally, the model may only be sensitive to certain parameters in a specific context. Consider a more generic version of the above rule (model parameterization 1), which hypothetically leads to biologically plausible model output:

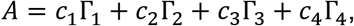

Which more completely, would be written:

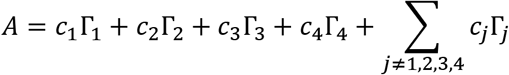

Where *c*_*j*_ = 0. There is no supposition that this hypothetically calibrated rule is uniquely correct. Assume the existence of an additional/alternative calibrated rule (model parameterization 2), which leads to equally biologically plausible model output:

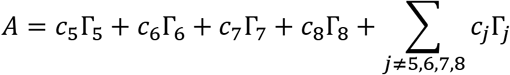

In a model parameterization 1, in which all cytokine multipliers are set to 0 except for cytokines 1-4, the model will appear more sensitive to cytokines 1-4 than the others. In model parameterization 2, the model will appear more sensitive to cytokines 5-8. A sensitivity analysis on one model parameterization is not valid for all other model parameterizations. A comprehensive sensitivity analysis would then have to incorporate the model configuration under which it is being performed, which is computationally intractable for all but the smallest of models.

We present this work as an alternative to the traditional model calibration/sensitivity analysis process described above. We blend the two distinct development stages into a single calibration stage with the end goal being a parameterization that matches the data and its associated variance throughout time. To begin, we posit that there are two primary factors responsible for biological heterogeneity in experimental data sets: stochasticity and genetic/epigenetic variation among individuals, the latter of which is manifest a variable responsiveness of the various pathways that determine cellular behavior. It is well known in biology that the systemic response to identical perturbations in genetically identical individuals (i.e., mice) is governed according to some probability distribution. This small stochastic variability in response can propagate over time such that it ultimately leads to divergent phenotypes. As such, ABM’s must incorporate some degree of randomness to simulate these behaviors. However, solely incorporating stochasticity into model rules is insufficient to capture the full range of bio-plausible model output – genetic/epigenetic variation among *in silico* test subjects must also be represented. The *in-silico* analogue to the human genome/epigenetic state is the specific parameterization of an ABM’s rule set. In order to represent a biological population, there must exist a range on each parameter within the rule-set parameterization. Here we re-emphasize our assertion that the parameters for the expressed ABM rules represent a compilation of “hidden” pathways that govern the responsiveness of the explicitly represented pathways; thus, the parameters represent the combination of an individual’s genetic and epigenetic makeup that affect the function of those “hidden” pathways.

In order to demonstrate this, we utilize a previously developed an ABM of systemic inflammation, the IIRABM. Though the IIRABM has been calibrated to simulate blunt trauma and infectious insult, it is an abstract and generalizable (9, 28) model of human response to injury. Cytokine time series and systemic response varies significantly between both blunt trauma/infectious insult and severe/large surface area burns. In this work, we demonstrate the changes necessary to recalibrate the model from simulating an infectious injury to a caustic and sterile injury. The IIRABM is a two-dimensional abstract representation of the human endothelial-blood interface. This abstraction is designed to model the endothelial-blood interface for a traumatic (in the medical sense) injury and does so by representing this interface as the unwrapped internal vascular surface of a 2D projection of the terminus for a branch of the arterial vascular network. The closed circulatory surface can be represented as a torus, and this two-dimensional surface defines the interaction space simulated by the model. The spatial geometry of the circulatory system and associated organ interfaces are not directly mapped using this scheme. This abstraction reproduces the circulatory topology accessible by the innate immune system and presents a unified means of representing interaction between leukocytes and endothelial surfaces across multiple tissue and organ types. The IIRABM utilizes this abstraction to simulate the human inflammatory signaling network response to injury; the model has been calibrated such that it reproduces the general clinical trajectories of sepsis. The IIRABM operates by simulating multiple cell types and their interactions, including endothelial cells, macrophages, neutrophils, T-lymphocyte subtypes (TH0, TH1, and TH2 cells) as well as their associated precursor cells. The simulated system dies when total damage (defined as aggregate endothelial cell damage) exceeds 80%; this threshold represents the ability of current medical technologies to keep patients alive (i.e., through mechanical organ support) in conditions that previously would have been lethal. The IIRABM is initiated using 5 parameters representing the size and nature of the injury/infection as well as a metric of the host’s resilience: 1) initial injury size, 2) microbial invasiveness (rate at which infection spreads), 3) microbial toxigenesis (rate at which infection damages tissue), 4) environmental toxicity (amount of spontaneous infectious exposure in the environment, such as an Intensive Care Unit), and 5) host resilience (the rate at which damaged but not dead tissue recovers). These 5 parameters clearly have correlates in the real world yet are nearly inherently un-quantifiable. Therefore, they are treated as dimension-less coordinate axes in which the behavior of the IIRABM exists.

The IIRABM characterizes the human innate immune response through the generation of a suite of biomarkers, including the pro-inflammatory and anti-inflammatory cytokines included in the IIRABM. At each time step, the IIRABM measures the total amount of cytokine present for all mediators in the model across the entire simulation. The ordered set of these cytokine values creates a high-dimensional trajectory through cytokine space that lasts throughout the duration of the simulation (until the *in silico* patient heals completely or dies). We note that stochastic effects can play a significant role in simulation dynamics. Model parameterizations used in this work lead to a simulated mortality rate of 50%; in these simulations, identical injuries and initial conditions are given to the model and over time, the trajectories diverge to the point that half of the simulated cohort heals completely and half dies. The fact that the initial conditions are exactly identical means that it is indeed stochasticity, not chaos, that leads to the diverging trajectories. A detailed discussion of this can be found in (9).

While the IIRABM successfully simulates the human immune response to injury at a high level (outcome proportions, time to outcome, etc.), it may not always replicate specific cytokine time series. A cytokine time series is not a single sequence of numerical values; rather, it is a sequence of ranges, indicating significant heterogeneity clinical response to severe burns, within which the cytokine measurements fall for a given patient in the cohort that generated the time series. This is challenging because the magnitude of these ranges is not temporally constant. In order for a computational model to be biologically realistic, it must be able to generate any physiological state which can experienced by the biology which is being simulated, and do so with the appropriate frequency – we posit that some small/simple set of stochastic differential equations with an analytically defined “noise” term cannot hope to explain nontrivial clinical data. In this work, we use Genetic Algorithms (GA) to operate on the IIRABM’s rule set such that it can accurately simulate the cytokine time course and final outcomes for a serious burn injury. Cytokine time series were extracted via inspection from (29). In (29), Bergquist, et al, provide a variety of blood cytokine levels over 15 time points and 22 days for patients which exhibited severe burns over 50% of the surface area of their bodies. The authors observed a mortality rate of 50% for this category of injury.

## Results

As a first attempt at optimization, we weighted the contributions of each of the five cytokines equally. We obtained excelled results for four out of five of the comparison cytokines, however the model was not optimized well to generate IL-10 concentrations which matched the literature, with peaking occurring at 6 hours post-insult rather than 5 days post-insult, as was seen clinically Figure 1A). We also posit that the nature of the IL-10 time series makes a poor fit more likely when using a fitness function that weights the contributions of each cytokine equally; the IL-10 time series data spikes at t-5 days but is near zero everywhere else. A candidate MRM parameterization that minimizes IL-10 production over the entire time course would then contribute less to the overall fitness (in this case, we seek to minimize the fitness function) than a hypothetical parameterization that was 10% off on TNF levels for every time step. In order to address this, we both doubled and tripled the weight of the IL-10 contribution to the fitness function; both of these modifications showed similar improvements over the initial fitness function, but neither was significantly better than the other. This leads us to expect that a doubling of the IL-10 contribution to the fitness is sufficient. We display this difference in Figure 1.

**Figure 1:**
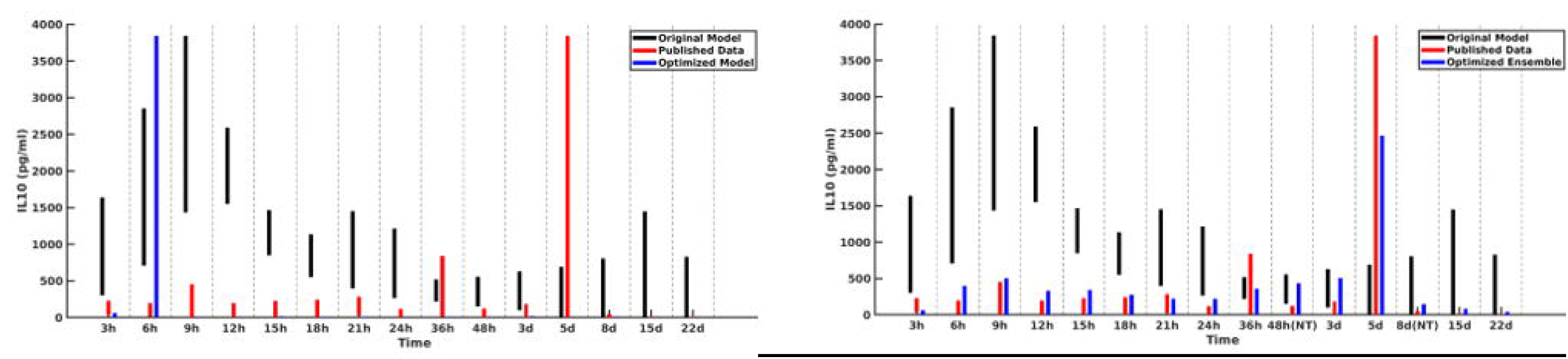
Cytokine ranges are shown for IL-10 for the original model (black), published data (red), and optimized ensemble model (blue). On the left Panel A, the IL-10 contribution to the fitness function is weighted equally to the other cytokines, with the result that simulated IL-10 levels after 6 hrs are essentially 0; on the right Panel B, the IL-10 contribution to the fitness function was doubled, and the cytokine dynamics more closely mirror those seen clinically.

A plot of cytokine ranges for 5 cytokines which existed in the clinical data set and were already present in the model at the start of this work (GCSF, TNF-α, IL-4, IL-10, and IFN-γ) is shown in Figure 2. Ranges for the original model, described in (9, 10), are shown in black; ranges for the published data (29) are shown in red;; and results from the optimized ensemble model are shown in green.

**Figure 2:**
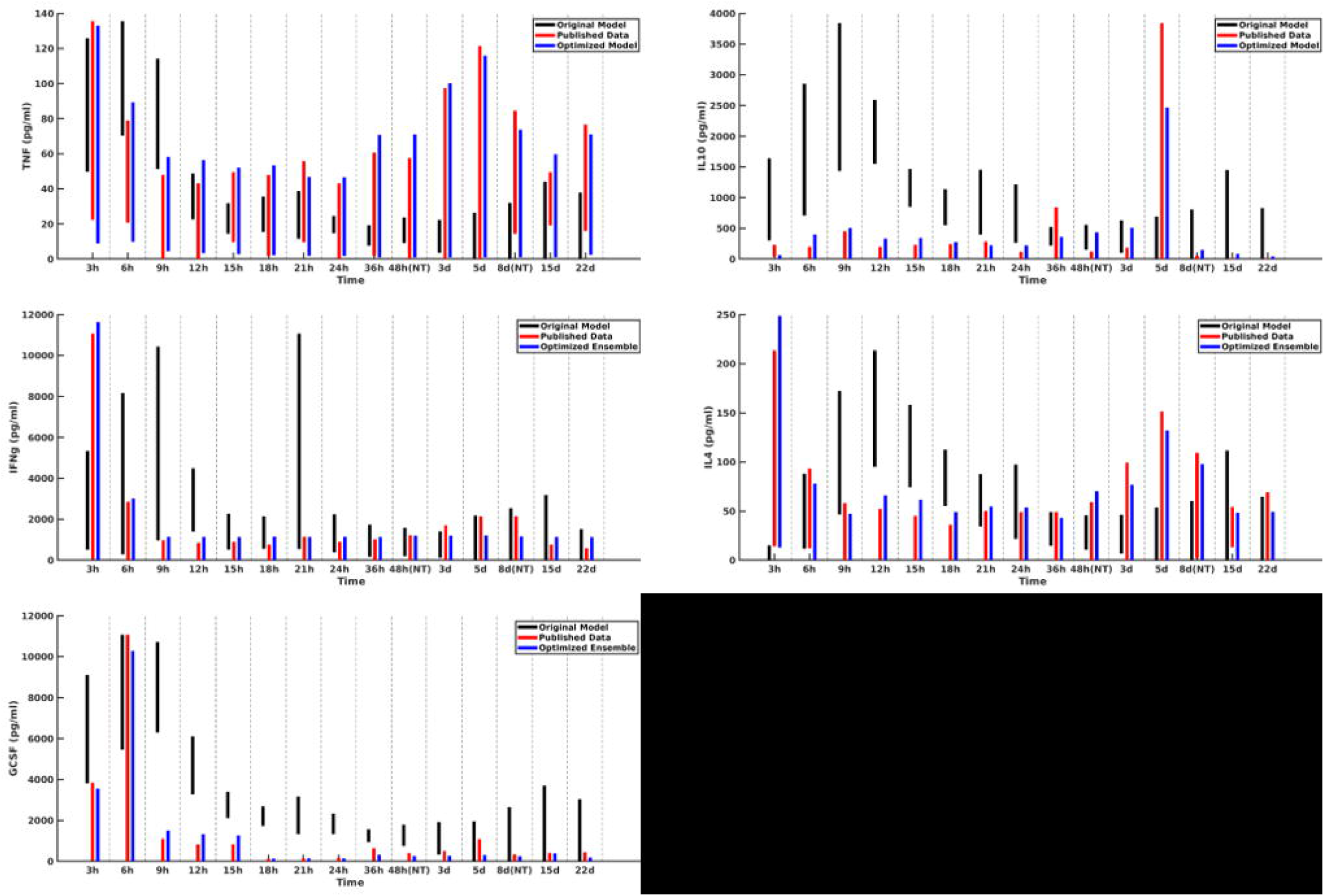
Cytokine ranges are shown for the original model (black), published data (red), and optimized ensemble model (blue) for TNFα (top-left), IL-10 (top-right), IFNγ (center-left), IL-4 (center-right) and GCSF (bottom-left). Ranges for the computational models were generated using 50 stochastic replicates.

The temporal cytokine dynamics expressed by the IIRABM are significantly modified from its original incarnation. We note that the ensemble model is optimized to match four out of five of the cytokines used in the fitness function to be nearly indistinguishable from the clinical data. We note a slight under-expression of IL-10 at t=5 days post injury. This discrepancy identifies a weakness in our model when it is being used to simulate burns, namely, that the cellular production of IL-10 is not well enough defined, in that its production is limited to activated macrophages and TH2 helper cells. Given that the IIRABM was developed to represent the innate immune response to traumatic injury, we consider this recalibration to burn injuries to be a success.

In Fig. 3, we compare the original rule matrix to the optimized rule matrix. Numerical values for both matrices can be found in the supplemental material. The optimized matrix has a much more connected structure, and is a dense matrix, as opposed to the sparse original rule matrix. There are not any matrix elements with a value of 0 in the optimized matrix, though there are many elements with comparatively small values. This structure is similar to what is seen in experimental bioinformatic studies; all of the cytokines in this network appear to be connected to each other, at least to a small degree, while a smaller number of strong connections (which could also be considered correlations) provide the majority of the influence on the system dynamics. The original rule matrix, with complete labeling, can be found in the supplementary material.

**Figure 3:**
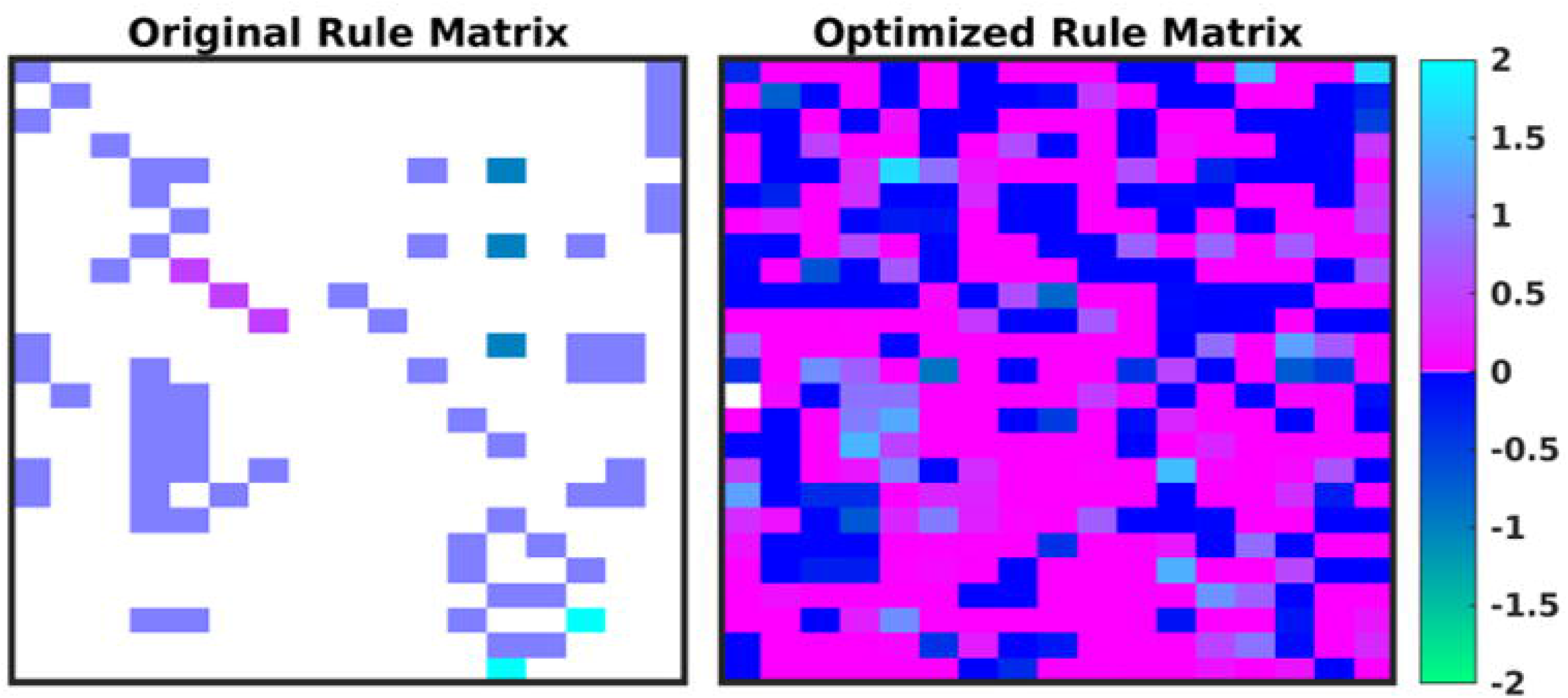
A heatmap of the original rule matrix is shown on the left and the optimized matrix is shown on the right. In these heatmaps, the white blocks represent a matrix element with a value of 0 (e.g. no connection); the dark blue to green represents a negative matrix element; the pink to light blue represents a positive matrix element. Optimization vastly increases the connectivity of the ABM elements (as would be expected in the true biological system). Rule definitions and labels are found in the supplementary material.

In Fig. 4, we present the time evolution of the diversity of the simulated population. We define the total diversity of a population to be the sum of the ranges of each matrix element. In Panel A, matrix element ranges are ordered from low to high. In the first several generations, diversity is maximized over the entire matrix. As the system evolves towards an optimum parameterization, diversity decreases, and the matrix begins to converge to a single value. In order to combat this, we use a mutation rate that increases as a function of the generation number, which begins to reintroduce diversity into the population. This is seen in Panel A, as the matrix element ranges begin to return to a diverse configuration, and more globally in Panel B, which plots the total diversity metric as a function of generation number.

**Figure 4:**
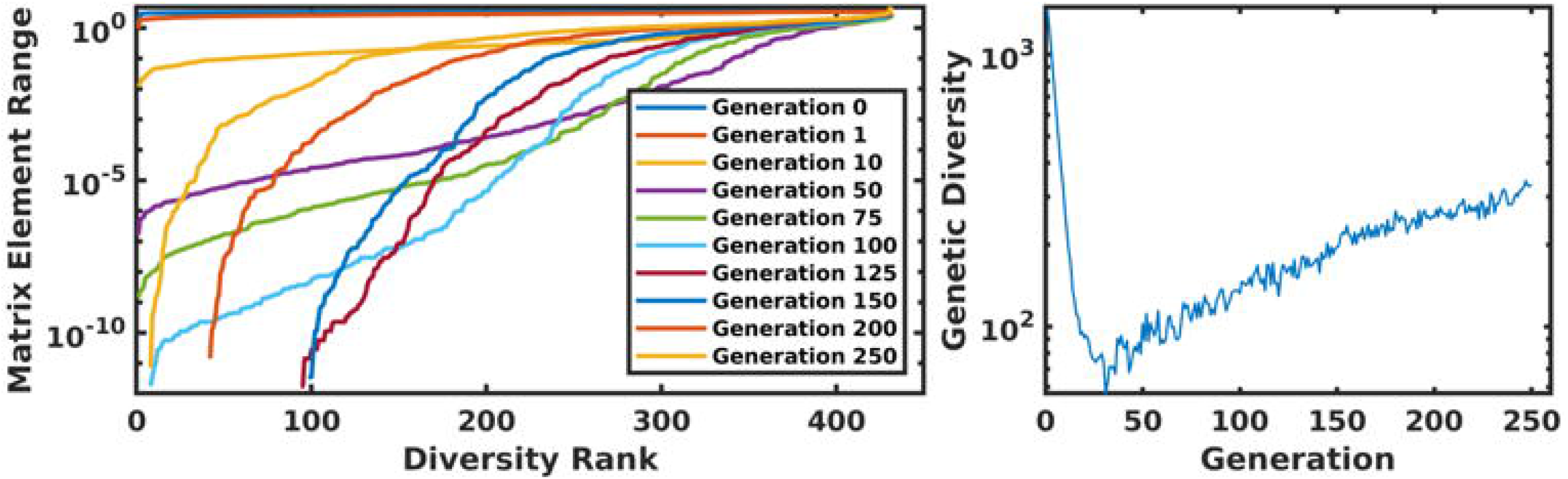
Panel A displays the ordered matrix element ranges for a variety of time points throughout the genetic algorithm. In this plot, the most diverse generations are represented by a nearly-horizontal line at the top of the plot. As the system evolved, this diversity begins to collapse until the increasing mutation rate compensates for the algorithm’s convergence. This is displayed in Panel B, which shows the total diversity of the population as a function of generation number.

## Discussion

The IIRABM rule set utilized in this work contained 432 free and continuous parameters, many of which had highly nonlinear or conditional effects on the model-generate cytokine trajectories and outcomes. Due to cytokine-specific properties, IL-10 was more challenging than the others when performing a multi-cytokine time series optimization. We acknowledge that model validation is a crucial component of the modeling process and do not contend that this work provides a strong validation of the IIRABM as a model of burns. Ultimately, the model can only be as strong as the data used to inform it. Rather, the subject of this paper is to present an alternative means of calibrating a computational model to a data set while maintaining the capability to represent the heterogeneity of the data, thereby potentially reflecting critical biological processes that account for the ubiquitous inter-individual variability seen in biological systems. This raises the critical issue of fitness function design in terms of how the calibration process proceeds.

We note that if we were to set the fitness function to match the published data as closely as possible we would be limiting the representational capacity and potential generalizability of the model, as the true range of biologically plausible blood cytokine concentrations in undoubtedly larger than what is seen in a cohort of 20 patients (this is exactly the Denominator Problem as described in the Reference in Bulletin of Mathematical Biology (27)). In order to obtain a more generalizable model, we propose two alternatives approaches to the above presented work: 1) that the fitness function should be configured to over-encompass the available data, with cytokine range boundaries determined by the probability density function (pdf) which governs the experimental data; or 2) synthesize multiple datasets in order to design a fitness with maximum cytokine rage coverage that is still supported by experimental data. Incorporating the shape of the probability density function into the fitness function can be difficult purely as a matter of practicality – often the raw data for human cytokine levels isn’t available, and only the absolute range can be extracted from published manuscripts, and it is also common to see a cohort size that is too small to definitively propose a single pdf which adequately describes the data.

In future work, we will utilize this genetically diverse *in silico* cohort as part of our machine-learning therapeutic discovery workflow (10, 30). We note the importance of *in silico* genetic diversity for therapeutic discovery in (10); in this work, we developed a multi-cytokine/multi-time-point therapeutic regimen which decreased the mortality rate from ~80% to ~20% for a severe simulated injury. The therapy was discovered using genetic algorithms on a single model internal parameterization. When we examined the non-responders, we noted that hyperactivity in specific pathways could manifest negatively, specifically, excess GCSF activity lead to excess neutrophil recruitment, which instigated a state of perpetual inflammation. Brittle policies/solutions (i.e., those that are not applicable outside of the very specific circumstances used to train them) have long been recognized as a weakness of machine learning research (31). In order to overcome this obstacle, data used to train machine-learning algorithms should be sourced as broadly as possible. A useful analogy would be to compare the machine learning experiment to an *in vivo* biological experiment: performing a biological experiment on a set of genetically identical animals will yield less generalizable information than an experiment performed on a set of genetically heterogenous animals.

Further, we note that, while we generated a diverse *in silico* patient cohort which generates cytokine trajectories that match clinical data, the diversity is limited by the algorithm. A GA is an optimization algorithm, meaning that the algorithm attempts to find a path through parameter space which leads to some optimum of the fitness function. Thus, even though we collect viable parameterizations as the algorithm progresses, they are sampled from a limited region of parameter space. Future work will seek to more comprehensively explore the entire parameter space using active learning, similar to (32).

The result of GA model refinement and rule discovery is a new rule-set and associated parameterization, discovered through machine-learning, which, when instantiated dynamically, generates data that matches what is seen experimentally; in reality, this result represents a single step in an iterative cycle of model refinement and biological experimentation. At the conclusion of the GA run, there exists an ensemble of candidate model parameterizations which meet the fitness criterion to some fixed threshold. Many of the genes in each individual parameterization end up tightly constrained by the algorithm, while others have a larger range. These latter parameters are those about which the model is most uncertain. Active Learning is a sampling technique used in machine learning in which sampled data is chosen based on how much information it can apply to the machine learning model. A similar approach can be taken in this case. In order to most efficiently update and refine the computational model, experiments should be designed to query the model features that are most uncertain. This approach is illustrated in Fig. 5. In this way, GA can play an integral role in the iterative cycle of model refinement and experimentation necessary to construct a high-fidelity generalizable computational model.

**Figure 5:**
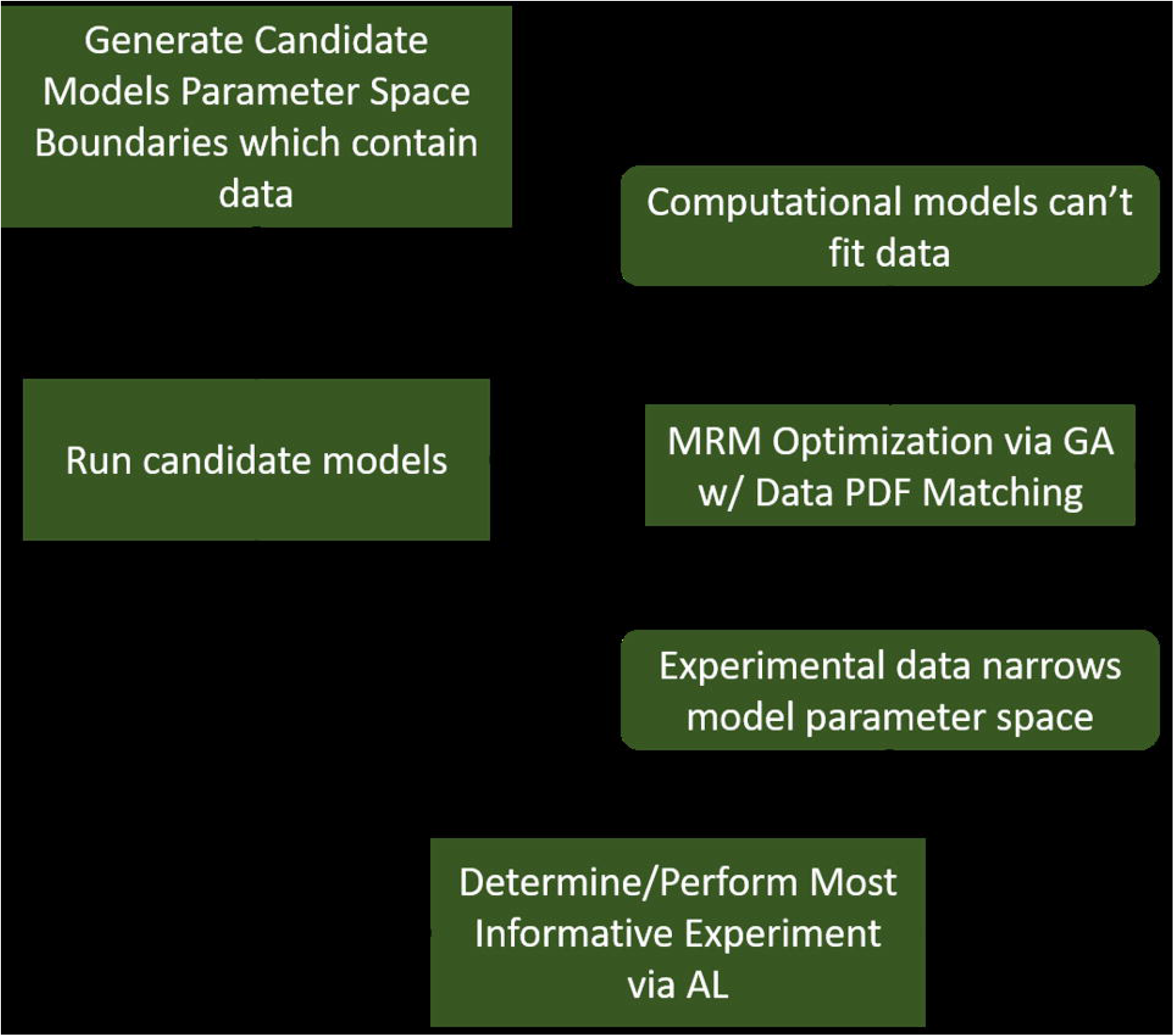
A diagram indicating a hybrid experimental/computational workflow for the automated calibration and validation of ABMs using the MRM scheme. In this workflow, a computational model containing all mechanistic knowledge hypothesized to be relevant to the biological system in question is developed. The range of output for a comprehensive set of viable model parameterizations is determined and compared to biological data. At this point, experimental data can be used to eliminate some of the formerly viable model parameterizations or invalidate the model. In the event the model is invalidated, it can be redesigned/reconfigured to address its shortcomings. After that, the remaining set of putative model parameterizations is investigated to determine which specific parameters contribute the most variability to the model output. These are then the parameters that are selected for further characterization in subsequent biological experiments.

## Conclusion

In this work, we have utilized genetic algorithms to operate on the rule structure of an existing agent-based model of the human innate immune response to a traumatic injury in order more comprehensively reflect the biological heterogeneity seen in the clinical setting. The inability of prior approaches to approximate the heterogeneity seen in experimental and clinical data sets remains a significant challenge in the use of complex multiscale models, such as ABMs, and the approach presented in this paper addresses the associated issues of tuning the model’s rule-set by either the addition of new rules or the re-parameterization of existing rules. In the presented example this has been accomplished by simulating the human innate immune response to a severe burn injury while reproducing the variation seen in a clinical burn data set. In order to mitigate the risks of overfitting, we have utilized the genetic algorithm as a parameter-space exploration tool, rather than a pure optimizer; in the process of optimization, any candidate solution that can explain the clinical data is retained. The end result is a diverse ensemble of putative model parameterizations, each of which generates cytokine time series and outcomes that fall within the clinical ranges seen in the data used to train the model. Ultimately, the synthesis of machine learning and high-performance computing renders the parameter-space exploration of a high-dimensional agent-based model to be computationally tractable and does so in a fashion that provides increased translational utility by capturing biological heterogeneity.

## Methods

A GA (33–35) is a population-based optimization algorithm that is inspired by biological evolution. In a GA, a candidate solution is represented by a synthetic ‘genome,’ which, for an individual, is typically a one-dimensional vector containing numerical values. Each individual in a genetic algorithm can undergo computational analogues to the biological processes of reproduction, mutation, and natural selection. In order to reproduce, two individual vectors are combined in a crossover operation, which combines the genetic information from two parents into their progeny.

Using this scheme, cytokines produced by a given cell type are held fixed, while the stimuli that lead to the production of that specific cytokine are allowed to vary. This maintains a distinction between the cell and tissue types represented in the model throughout the MRM evolution from the GA.

The candidate genomes which comprise the rule set are then tested against a fitness function which is simply the sum of cytokine range differences between the experimental data and the computational model:

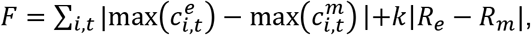

where 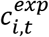 represents the normalized blood serum level of cytokine *i* at time point *t* from the experimental data, 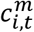represents the normalized blood serum level of cytokine *i* at time point *t* from the IIRABM, *R*_*e*_ represents the experimentally observed mortality rate, *R*_*m*_ represents the model-generated mortality rate, and *k* is an adjustable parameter to govern the importance of the mortality rate contribution to the fitness function. For the purposes of this work, we consider an optimal solution to be one that minimizes the above fitness function. In order to avoid issues of over-fitting, we held the time points at t=48 hrs. post-burn and t=8 days post-burn back from the evaluation of candidate fitness. Despite this, these time points were well-matched between the *in silico* and *in vivo* experiments.

Candidate genomes are then selected against each other in a tournament fashion, with a tournament size of 2 [28, 29]. The tournament winners make up the breeding pool, and progenitor genomes are randomely selected and paired. We implement a variant of elitism in that, at the completion of the tournament, the least fit 10% of the candidate progenitors are replaced with the fittest 10% of candidate genomes from the precious generation. Progeny genomes are defined with a uniform crossover operation using a standard continuous formulation (36):

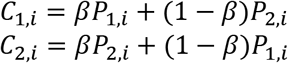

Where *C*_1,*i*_ is the value for gene *i* in child 1, *P* is the value for gene *i* in parent 1, and *β* is a random floating-point number between 0 and 1. After breeding, each child is subject to a random chance of mutation which begins at 1% and increases with each generation.

We employ an elitist strategy by replacing the least fit 10% of the breeding population with the most fit parameterizations. This ensures that our best solutions are not lost due to mutation. Additionally, we have used a nonviability criterion to filter out candidate parameterizations that we would not want to propagate. In this case, any parameterization which leads the model to die before the first clinical time point (3 hrs post injury) is discarded and replaced with a fitter candidate. This nonviability criterion is only triggered in the first few generations, as the algorithm quickly discards nonviable regions of parameter space.

In order to capture viable, but non-optimal model parameterizations, each candidate parameterization is checked to see whether or not cytokine ranges all stochastic replicates fall within the clinically-defined ranges. Those that generate trajactories that lie within the clinical boundaries are placed into the ensemble of biologically plausible models. Thus, the end result of this procedure is both an optimal model (that which fits the variance of the data best) and an ensemble of model parameterizations that generate data which is consistent with what is seen clinically.

The IIRABM was optimized for 250 generations with a starting population size of 1024 candidate parameterizations. The IIRABM was implemented in C++ and the GA was implemented in Python 3; and simulations were performed on the Cori Cray XC40 Supercomputer at the National Energy Research Scientific Computing Center and at the Vermont Advanced Computing Center. Codes can be found at https://github.com/An-Cockrell/IIRABM_MRM_GA. The final Python code used in this work is ga_wrapper_elitism_v6.py.

## Supporting Information Legend

The file, ‘InternalParameterization_H_RandCh_IS25_Gen249.csv’ is the full set of parameterizations for the 250^th^ generation/iteration of the genetic algorithm. This information was used to inform Figures 2 and 3. The file, ‘RuleMat.xslx’ is the rule matrix for the original version of the IIRABM. Rules are represented in the rows of the matrix and the contribution of each cytokine is represented in the columns of the matrix.

## List of Abbreviations

ABM: Agent-Based Model
GA: Genetic Algorithm
IIRABM: Innate Immune Response Agent-Based Model
MRM: Model Rule Matrix
HPC: High-Performance Computing
TNFα: Tumor Necrosis Factor alpha
IL-10: Interleukin 10
IFN-γ: Interferon gamma
2D: Two Dimensional

## Declarations

### Ethics

All human data was obtained from published and publicly available sources; no approval was required.

### Consent for Publication

Not applicable

### Availability of Data

Codes used to generate data used in this study can be found at https://github.com/An-Cockrell/IIRABM_MRM_GA. The final Python code used in this work is ga_wrapper_elitism_v6.py.

### Authors’ Contributions

CC developed the codes, ran the simulations, analyzed the results, and drafted the manuscript. GA participated in the design of the study and the writing of the manuscript. All authors read and approved the final manuscript.

## Acknowledgements

Not Applicable

